# Test-statistic correlation and data-row correlation

**DOI:** 10.1101/759027

**Authors:** Bin Zhuo, Duo Jiang, Yanming Di

## Abstract

When a statistical test is repeatedly applied to rows of a data matrix—such as in differential-expression analysis of gene expression data, correlations among data rows will give rise to correlations among corresponding test statistic values. Correlations among test statistic values create many inferential challenges in false-discovery-rate control procedures, gene-set enrichment analysis, or other procedures aiming to summarize the collection of test results. To tackle these challenges, researchers sometimes will—explicitly or implicitly—use the correlations (e.g., as measured by the Pearson correlation coefficients) among the data rows to approximate the correlations among the corresponding test statistic values. We show that, however, such approximations are only valid under limited settings. We investigate the relationship between the correlation coefficient between a pair of test statistics (test-statistic correlation) and the correlation coefficient between the two corresponding data rows (data-row correlation). We derive an analytical formula for the test-statistic correlation as a function of the data-row correlation for a general class of test statistics: in particular, two-sample *t*-test is a special case. The analytical formula implies that the test-statistic correlation is generally weaker than the corresponding data-row correlation, and in general, the latter will not well approximate the former when the involved null hypotheses are false. We verify our analytical results through simulations.

## 1. Introduction

Many scientific data sets are organized in matrix forms and statistical inferences—such as hypothesis tests and regression analysis—are often repeatedly applied to individual rows of the data matrix. For example, in gene expression analysis, normalized expression values are often organized in a matrix with rows corresponding to genes and columns corresponding to biological samples (experimental units). In a two-group comparison experiment, a two-sample test will be applied to each row of the data matrix in order to assess differential expression (DE). For more complex experimental designs, regression analysis can be used.

Correlations may exist among the data rows: For example, between-gene correlations are commonly observed in gene expression data [1, 2, 3, 4, 5]. Data-row correlations can give rise to correlations among the test statistic values calculated from the data rows [6, 7, 8]. The dependence among test statistic values has brought methodological challenges to statistical procedures aiming to summarize the collection of test results. For example, some multiple hypothesis testing procedures determine a *p*-value cutoff by controlling the *false discovery rate* (FDR) [9] or the *q-value* [5]. Many FDR-control procedures are valid only when the test statistics satisfy certain independence or positive-dependence conditions [9, 10]. Furthermore, Efron [7] showed in a simulation study that for a nominal FDR of 0.1, the actual false discovery proportions (FDP) in individual experiments can easily vary by a factor of 10 when there are correlations among test statistics.

In a gene-set analysis, one tests for over-abundance of DE genes in a specified gene set (e.g., a molecular pathway or a gene ontology category) [11]. The correlations among DE test statistics, if not addressed appropriately, will undermine the validity of many gene-set tests [2, 8, 12]. A better understanding of the test-statistic correlations is thus of fundamental importance and is a first step towards developing statistical methods that correctly account for test-statistic correlations.

Without replicating the experiment, we cannot directly estimate the correlation between a pair of test statistic values, because there is only one observed test statistic value for each data row. For this reason, the correlation between the corresponding data rows (after treatment effects accounted for) is sometimes used as a surrogate—explicitly or implicitly—when one actually needs the test-statistic correlation. It is yet unclear when and to what extent the teststatistic correlation (e.g., as measured by the Pearson correlation coefficient) can be approximated by the corresponding data-row correlation, though some simulation results suggest connections between the two quantities. Efron [7] concluded through simulation that the distribution of *z*-value (the test statistic considered in that paper) correlation can be nearly represented by the distribution of sample correlation from the data rows. Barry et al. [6] showed by Monte Carlo simulation of gene expression data that a nearly linear relationship holds between test-statistic correlations and data-row correlations for several forms of test statistics they examined. These Monte Carlo simulation results were cited by Wu and Smyth [8] as a justification for estimating a variance inflation factor from data-row correlations in order to correct for test-statistic correlations.

In this paper, we derive an analytical formula for the test-statistic correlation as a function of the data-row correlation for a general class of test statistics—including the familiar two-sample *t*-test as a special case. We use simulation results to confirm our analytical findings. We show that 1) the test-statistic correlation is equal to data-row correlation when the test statistic is a linear combination of the observed data, but 2) in general, the test-statistic correlation is weaker than and not well approximated by the corresponding data-row correlation. In particular, our analytical formula reveal that 3) the test-statistic correlation depends on whether the test statistic has an expectation of 0 (which often corresponds to whether the null hypothesis is true). These findings urge us to give more thoughts about correlations when trying to summarize the collection of the test results.

## 2. Methods and Results

Suppose we have a data matrix and have applied a statistical test to individual rows of the data matrix. We will consider pairwise correlations and focus on two rows of the data matrix: ***X*** = (*X*_1_, …, *X*_*n*_)^*T*^ and ***Y*** = (*Y*_1_, …, *Y*_*n*_)^*T*^ with mean vectors ***μ***_*X*_ and ***μ***_*Y*_. We will assume that the columns of the data matrix are independent so that (*X*_*j*_, *Y*_*j*_), *j* = 1, …, *n*, are independent bivariate random variables: this assumption is usually reasonable in a designed experiment for two-group comparison. The mean of (*X*_*j*_, *Y*_*j*_) may vary across experimental units *j* = 1, …, *n*, but we assume that the population variance-covariance structure remains the same across experimental units, that is,

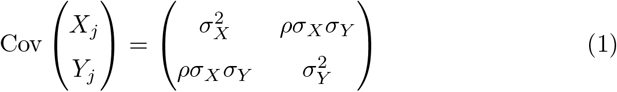

for all *j* = 1, …, *n*. We consider a general class of test statistic of the form

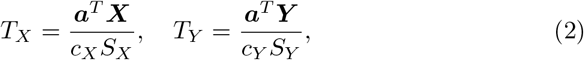

where ***a*** is a non-zero n-vector, (*c*_*X*_, *c*_*Y*_) are non-random constants, and (***a***^*T*^ ***X***, ***a***^*T*^ ***Y***) and (*S*_*X*_, *S*_*Y*_) are independent. In particular, the familiar two-sample *t*-test is of this form with *S*_*X*_ and *S*_*Y*_ estimating σ_*X*_ and σ_*Y*_ respectively. So is the *t*-test for a regression coefficient in a linear regression model.

We want to investigate the connections between the test-statistic correlation *ρ*_*T*_ = Cor(*T*_*X*_, *T*_*Y*_) and the data-row correlation *ρ* = Cor(*X*_*j*_, *Y*_*j*_) (common to all units *j*). First, we present an analytical formula that relates *ρ*_*T*_ to *ρ*.

### Theorem 1.

*For the test statistics T*_*X*_, *T*_*Y*_ *in (2):*

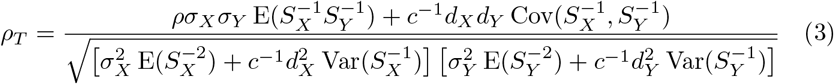

*where d*_*X*_ = ***a***^*T*^ ***μ***_*X*_, *d*_*Y*_ = ***a***^*T*^ ***μ***_*Y*_ *and c* = ***a***^*T*^ ***a***.

*Proof.* (*c*_*X*_, *c*_*Y*_) do not affect correction and can be ignored. For any (*U*_*X*_, *U*_*Y*_) that are independent of (*S*_*X*_, *S*_*Y*_), direct calculation shows that

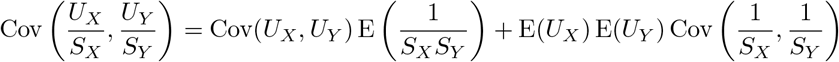

and

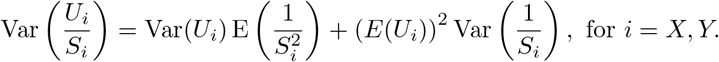

For this theorem, we let *U*_*X*_ = ***a***^*T*^ ***X***, *U*_*Y*_ = ***a***^*T*^ ***Y***, then E(*U*_*X*_) = ***a***^*T*^ ***μ***_*X*_, E(*U*_*Y*_) = ***a***^*T*^ ***μ***_*Y*_, 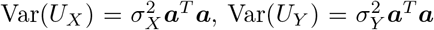, and Cov(*U*_*X*_, *U*_*Y*_) = *ρ*σ_*X*_ σ_*Y*_ ***a***^*T*^ ***a*** since the columns of the data matrix are assumed independent.

To apply equation (3) in practice, we need to compute the involved moments of 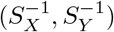, but equation (3) offers some insights without explicit calculation of those quantities.

### Corollary 1.

*ρ*_*T*_ = *ρ if S*_*X*_ *and S*_*Y*_ *are constants (i.e., not random)*.

*Proof.* When *S*_*X*_ and *S*_*Y*_ are constants, 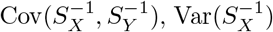 and 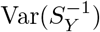 are all 0, and 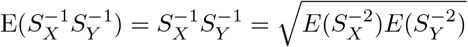.

This corollary says that for *z*-tests, the test-statistic correlation is the same as the corresponding data-row correlation ([6] also pointed out this). This confirms the simulation results in [7]. Another intuition offered by equation (3) is that the relation between *ρ*_*T*_ and *ρ* depends on whether one or both of ***a***^*T*^ ***μ***_*X*_ and ***a***^*T*^ ***μ***_*Y*_ are 0—which often corresponds to whether the corresponding null hypotheses are true. When both ***a***^*T*^ ***μ***_*X*_ and ***a***^*T*^ ***μ***_*Y*_ are 0, equation (3) will have a simpler form

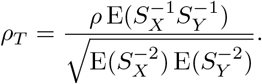

Intuitively, in such cases, we can expect *ρ*_*T*_ ≈ *ρ* in large samples if *S*_*X*_ and *S*_*Y*_ are “good” estimators of σ_*X*_. and 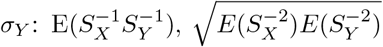 will then both tend to 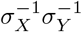.

More generally, though, the test-statistic correlation *ρ*_*T*_ is not the same as the data-row correlation *ρ*. Next, using the important special case of two-sample *t*-test, we will further demonstrate that, in general, *ρ*_*T*_ is not well approximated by *ρ*, even in large samples.

For equal-variance two-sample *t*-test, we let 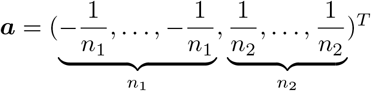,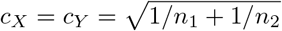 and 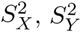 be the pooled sample variances,

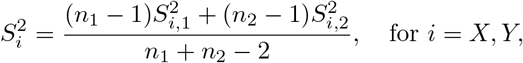

in (2), where 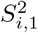 and 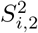 are the sample variances for sample 1 and sample 2 respectively in data row *i*. From Basu’s lemma, (***a***^*T*^ ***X***, ***a***^*T*^ ***Y***) are independent of (*S*_*X*_, *S*_*Y*_). Typically, the null hypotheses to test are *d*_*X*_ = ***aμ***_*X*_ = 0 and *d*_*Y*_ = ***aμ***_*Y*_ = 0.

### Theorem 2.

*For the equal-variance two-sample *t*-test, when n* = *n*_1_ + *n*_2_ → ∞ *and n*_1_/*n* → *r for some r*, 0 < *r* < 1,

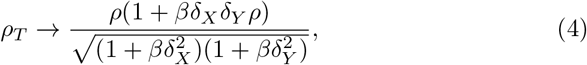

*where δ*_*X*_ = *d*_*X*_/σ_*X*_, *δ*_*Y*_ = *d*_*Y*_/σ_*Y*_, *and β* = *r*(1 − *r*)/2.

*Proof.* As *n* = *n*_1_ + *n*_2_ → ∞ and *n*_1_/*n* → *r*, in equation (3),

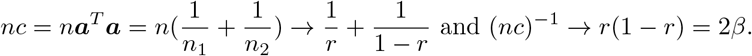

The key of the proof is to determine the limits of the moments 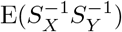,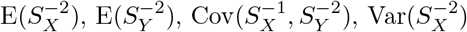, and 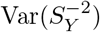. By the consistency of 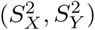 and the continuous mapping theorem,

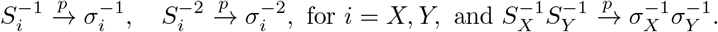

For large *v* (= *n* − 2), 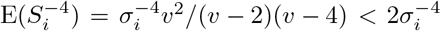, for *i* = *X, Y*. This implies the 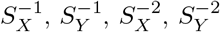 and 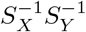 are all uniformly integrable (note that 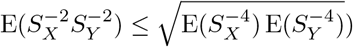, and thus

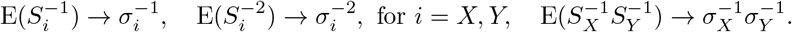

In the Appendix Lemma 1, we show that

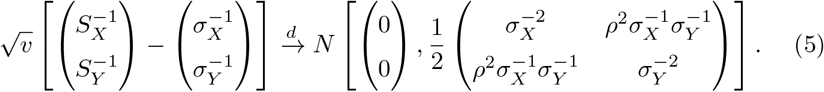

This together with the continuous mapping theorem suggests that

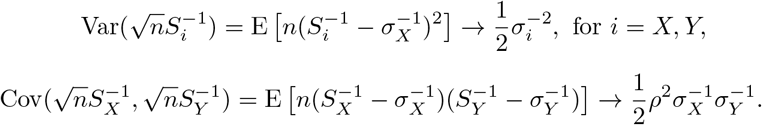

For these moments limits to hold, we need to show that the involved moments are uniformly integrable (see, e.g., Theorem 6.2 of [13]). It is sufficient to show that 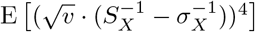 is bounded for large *v*: in the appendix Lemma 2, we show that

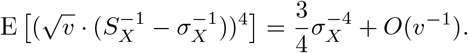

Plugging the limiting values of 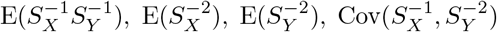, 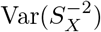, and 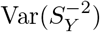 into equation (3) gives equation (4).

In the Appendix, we will also explain how to compute *ρ*_*T*_ in finite samples for two-sample *t*-test. It is mainly 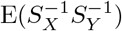 that is difficult to compute.

Theorem 2 (and equation (10) in the Appendix) reaffirms that the relation between *ρ*_*T*_ and *ρ* depends on (*δ*_*X*_, *δ*_*Y*_) = (*d*_*X*_/σ_*X*_, *d*_*Y*_/σ_*Y*_). Figure 1 shows the contour plot of the limiting value of *ρ*_*T*_ when *n*_1_ = *n*_2_ → ∞ (*r* = 1/2, *β* = 1/8) as a function of (*δ*_*X*_, *δ*_*Y*_), for *ρ* = −0.7, −0.1, 0.1, 0.7. Note that *ρ*_*T*_ → *ρ* if *d*_*X*_ = *d*_*Y*_ = 0: typically, this means both null hypotheses are true. One can show that | lim_*n*→∞_ *ρ*_*T*_ | ≤ |*ρ*|. That is to say, in general, the test-statistic correlation is weaker than the corresponding data-row correlation.

**Figure 1:**
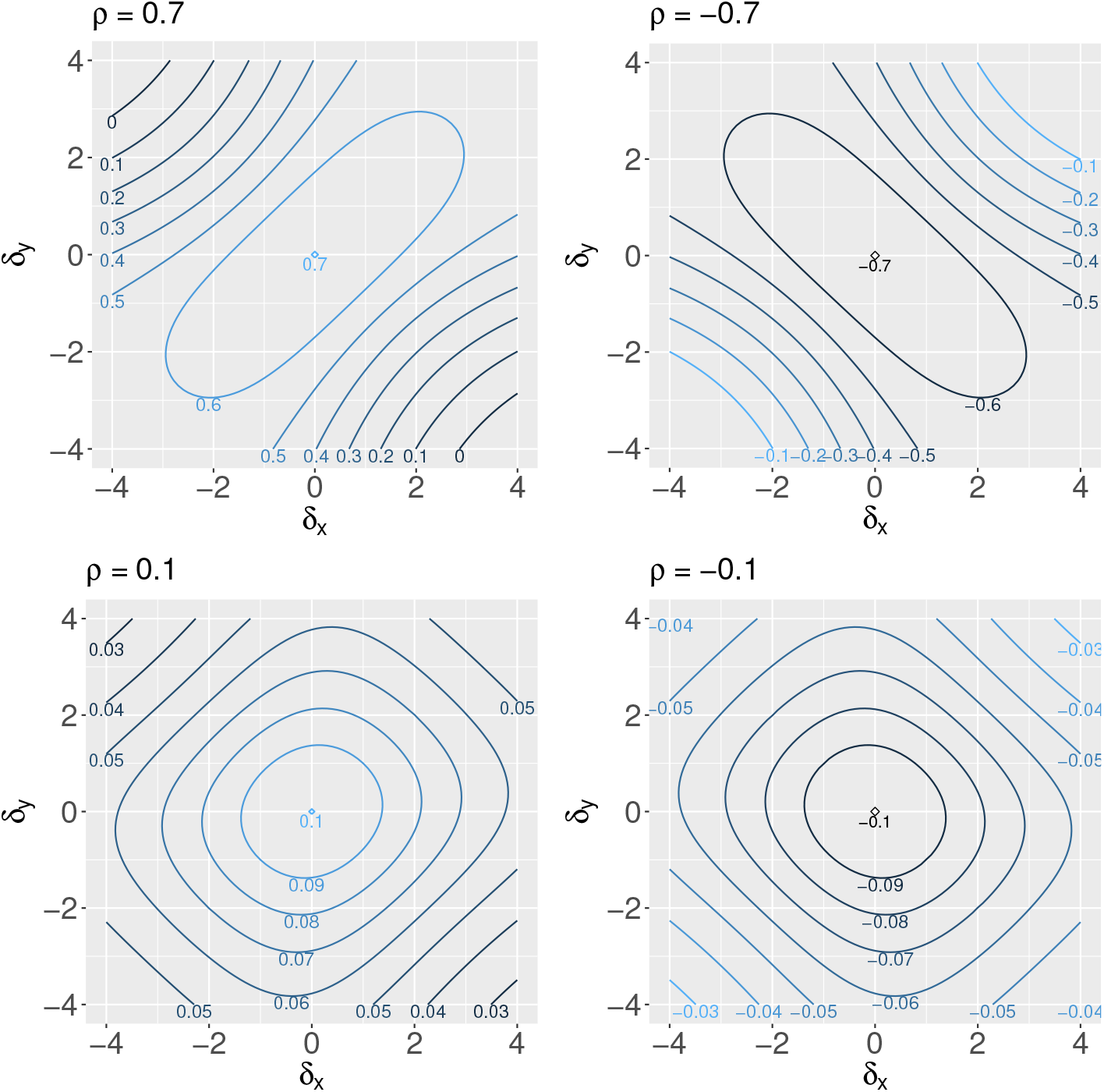
Contour plot of the limiting values of *ρ*_*T*_ as *n*_1_ = *n*_2_ → ∞. For each *ρ*, the asymptotic value of *ρ*_*T*_ is plotted as a function of (*δ*_*X*_, *δ*_*Y*_).

In Figure 2, we plotted *ρ*_*T*_ as a function of *ρ* when *n*_1_ = *n*_2_ = 3, 10 or ∞ for a few selected values of (*δ*_*X*_, *δ*_*Y*_). We also added simulated values of *ρ*_*T*_ (for *n*_1_ = *n*_2_ = 3, 10) to confirm our analytical findings: For each (*δ*_*X*_, *δ*_*Y*_) value, we let *ρ* vary form −1 to 1 by a step size of 0.01. For each *ρ*, we simulated a pair of data rows ***X***, ***Y*** according to independent bivariate normal distributions:

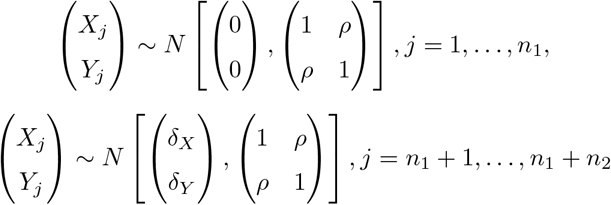

and computed the two-sample *t*-test statistics *T*_*X*_ and *T*_*Y*_. *H* = 5000 pairs of (*T*_*X*_, *T*_*Y*_) were simulated and their sample correlation 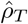 were shown in Figure 2.

**Figure 2:**
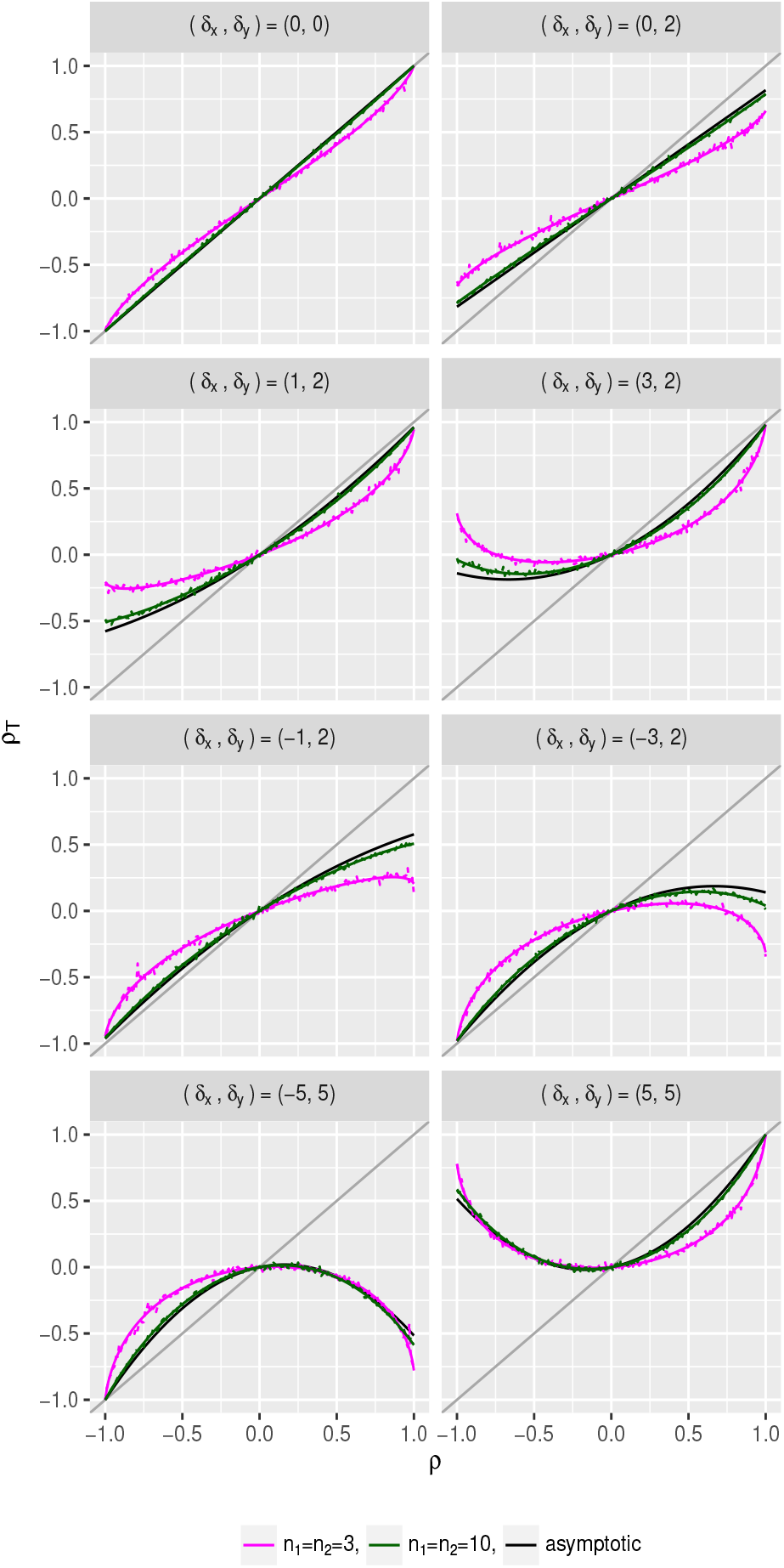
Test-statistic correlation *ρ*_*T*_ versus data-row correlation *ρ* at different (*δ*_*X*_, *δ*_*Y*_) values, when *n*_1_ = *n*_2_ = 3, 10, or ∞. The simulated values of *ρ*_*T*_ are also shown for *n*_1_ = *n*_2_ = 3 and 10. The solid (smooth) lines represent theoretical value, and dashed (jagged) lines represent simulated values.

Let 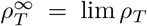 as *n*_1_ = *n*_2_ → ∞. We see from Figure 2 that when *δ*_*X*_ = *δ*_*Y*_ = 0, 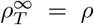; when *δ*_*X*_ = 0, 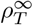 is a linear function of *ρ*; and when *δ*_*X*_ and *δ*_*Y*_ are both non-zero, 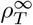 is a quadratic function of *ρ*. These features are predictable from the analytical formula (4) in Theorem 2 and they hold approximately in finite samples if *n* is large. In fact, we see that when *n*_1_ = *n*_2_ = 10, the *ρ*_*T*_ values are already remarkably close to 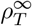.

In small samples (e.g., *n*_1_ = *n*_2_ = 3), there is more difference between *ρ*_*T*_ and 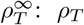 is often weaker than 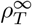 (i.e., 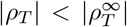) with a couple of exceptions (e.g., when *δ*_*X*_ = ±5, *δ*_*Y*_ = 5), which is reasonable since 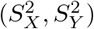 are “noisier” in small samples and noise in general reduces correlation. When both *δ*_*X*_ and *δ*_*Y*_ are non-zero (this typically means both null hypotheses are false), *ρ* does not approximate *ρ*_*T*_ well no matter what the sample size is: |*ρ*| can significantly overestimate |*ρ*_*T*_|. In extreme cases when *δ*_*X*_ *δ*_*Y*_ is big, *ρ*_*T*_ and *ρ* can have opposite signs.

## 3. Conclusion and discussion

This article discusses the relation between test-statistic correlation *ρ*_*T*_ and the corresponding data-row correlation *ρ*. Our results indicate that only in limited settings, *ρ*_*T*_ can be well approximated by *ρ*: for example, *ρ*_*T*_ = *ρ* for *z*-test and *ρ*_*T*_ ≈ *ρ* in large samples if both null hypotheses are true. For two-sample *t*-test, the relation between *ρ*_*T*_ and *ρ* will depend on (*δ*_*X*_, *δ*_*Y*_), the expected mean differences divided by the respective standard deviations of the data rows. When *δ*_*X*_ and *δ*_*Y*_ are both non-zero, *ρ*_*T*_ is a quadratic function of *ρ*, *ρ*_*T*_ can be much weaker than *ρ* (|*ρ*_*T*_ | < |*ρ*|), and *ρ*_*T*_ and *ρ* can sometimes have opposite signs.

Our findings have practical implications in statistical inferences aiming to summarize the collection of test results. For example, our results indicate that it is not reliable to approximate the distribution of test-statistic correlations by the distribution of data-row correlations if we expect the null hypotheses to be false for a significant proportion of the rows—which is often the case in gene expression analysis. If one wants to assess the null distribution of the test-statistic p-values by permuting the columns of the data matrix, then one has to realize the permutation will also change the correlations among the test-statistic values (since (*δ*_*X*_, *δ*_*Y*_) values will change after each permutation). In separate ongoing work, we are delving into these and related issues to better understand the impact of test-statistic correlation on gene set enrichment analysis, where one wants to test for overabundance of DE genes in a pre-specified set ([14] is one such attempt).

[8] discussed a variance inflation factor (VIF) which is useful when estimating the variance of the sum or average of m test statistics *t*_1_, *t*_2_, …, *t*_*m*_ when the corresponding genes (data rows) are correlated. In that paper, VIF is defined as 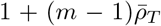, where 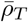 is the average of test-statistic correlations (i.e., *ρ*_*T*_ ’s) over all pairs of data rows in the set. (If all *t*_*i*_’s have the same variance *τ*^2^, then 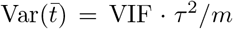) It was mentioned that 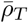 can be estimated by the average of data-row correlations. Our results indicate that replacing test-statistic correlations by data-row correlations will not be accurate if there are mean differences between the two groups among the data rows. For example, if we consider the two-sample *t*-test performed on *m* = 21 data rows in a matrix with correlated data rows (*ρ* = 0.1 for all pairs, variance σ^2^ = 1 for all rows) and mean differences ranging from −3 to 3 (uniformly spaced, i.e., *δ* = −3, −2.4, −1.8, …, 3 for the 21 rows) between two groups (*n*_1_ = *n*_2_ = 30), then the true VIF value computed using test-statistic correlations (which we can compute using asymptotic formula (4) in Theorem 2) should be 2.48; the VIF computed using the data-row correlations is 3.00, which overestimates the true VIF by 21%. In practice, we can estimate *ρ*_*T*_ for each pair of data rows by plugging the corresponding estimated values of *ρ, d, σ* values into equation (4). In the Appendix, we use a simulation to show that estimating VIF using estimated *ρ*_*T*_ values will outperform approximating *ρ*_*T*_ by the sample data-row correlations.

One reviewer asked whether our results apply to the moderated *t*-test where the variance estimation is based on a shrinkage method. The short answer is “no”. It is difficult to derive an analytical formula for the correlation between a pair of moderated *t*-test statistic values where a shrinkage method is used for estimating the variances of the test-statistic values, since information from all data rows are used for estimating the variances. Through a simple simulation where we applied moderated *t*-test in the `limma` package ([15]), we observed that the test-statistic correlations among moderated *t*-test statistic values still depend on *δ*_*X*_ and *δ*_*Y*_, but the relationship between test-statistic values and data-row correlations do not follow the analytical formula that we derived in Theorem 2. In particular, we observed that for moderated *t*-test, the teststatistic correlations tend to be greater than the data-row correlations in some cases where *δ*_*X*_ = *δ*_*Y*_ ≠ 0. For the usual two-sample *t*-test statistic, we have shown earlier that the magnitude of test-statistic correlation tends to be less than that of the data-row correlation. We included the details on the simulation settings and results of this simulation on moderated *t*-test in the Appendix.

In this paper, we assumed that the columns of the data matrix are independent and the explicit formula mainly focused on the two-sample *t*-test. We believe these are good starting points for discussing this complex issue. In the future, we plan to extend our investigation into more general settings: for example, the test for regression coefficients in a generalized linear model.

The R codes for reproducing the results in this paper are available at Github: https://github.com/zhuob/CorrelatedTest.

## Acknowledgement

Research reported in this article was partially supported by the National Institute of General Medical Sciences of the National Institutes of Health under Award Number R01GM104977 (to YD). We thank Sarah Emerson for valuable comments and suggestions in method development and manuscript preparation. This research was part of doctor dissertation of BZ under the supervision of YD. We thank the reviewers for their helpful comments.

## Appendix Lemmas for Theorem 2

We prove two lemmas for Theorem 2 under the theorem’s conditions and specifications:

### Lemma 1.

*Asymptotic distribution of* 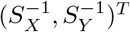

*Proof.* For the pooled variances in the two data rows ***X*** and ***Y***,

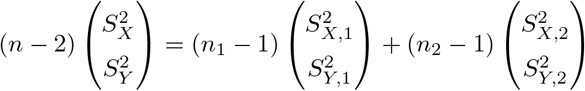

Let

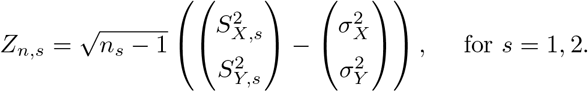

Then

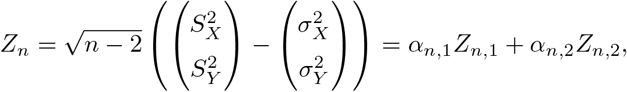

where 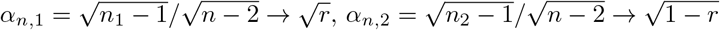.

From the central limit theorem (or the property of MLE), as *n*_1_, *n*_2_ → ∞, *Z*_*n*,1_ and *Z*_*n*,2_ are both asymptotically normally distributed as

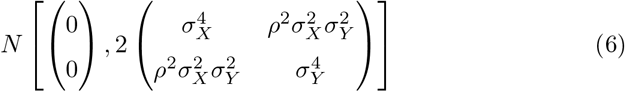

*Z*_*n*,1_’s and *Z*_*n*,2_’s are independent. So *Z*_*n*_ = *α*_*n,*1_*Z*_*n,*1_ + *α*_*n,*2_*Z*_*n,*2_ converges to the same distribution in (6) by Slutsky’s theorem and the continuous mapping theorem (note that 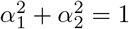).

Letting 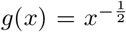 and applying *δ*-method to the asymptotic distribution (6) of *Z*_*n*_, we obtain (let *v* = n − 2)

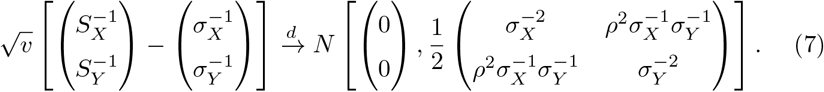

The next lemma shows that 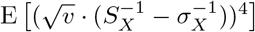 is bounded for large *v*.

### Lemma 2.

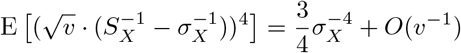

*Proof.* For the pooled sample variance, 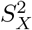, in the equal-variance two-sample *t*-test, 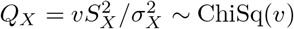 with *v* = *n* − 2 degrees of freedom, and

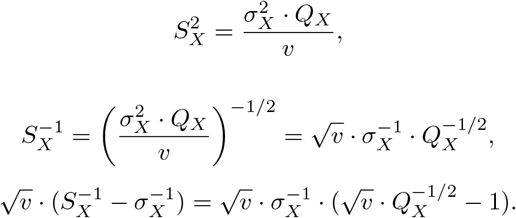

For a chi-square random variable with *v* degrees of freedom, *Q* ~ ChiSq(*v*),

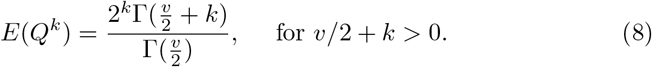

[16] (see also [17]) showed that for large *v*,

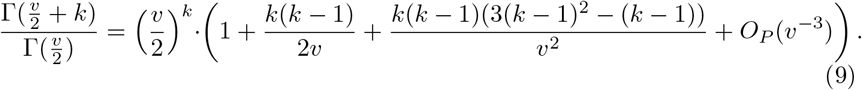

Letting *k* = −1/2, −1, −3/2, and −2, it then follows from (8) and (9) that

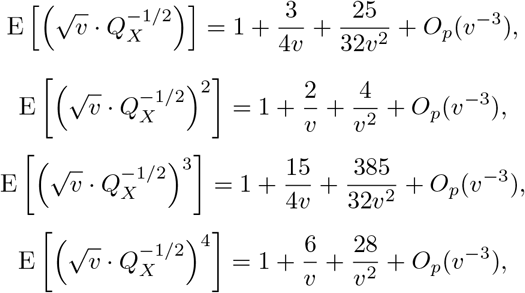

and thus

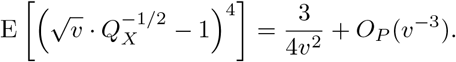

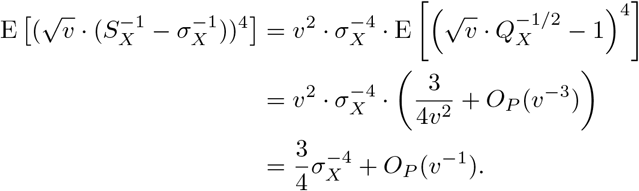

## Compute *ρ*_*T*_ in finite samples

It follows from (8) that for *v* > 2,

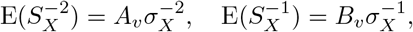

where

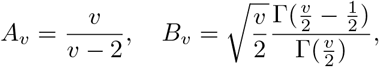

and thus

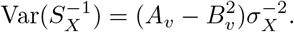

For bivariate normal data, [18] derived a formula (Theorem 3.1 in that paper) for computing product moments of the form 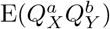 as infinite sums, for 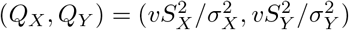:

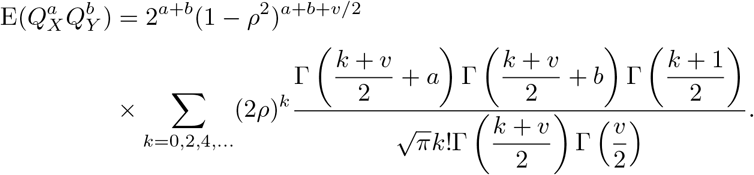

So we can numerically compute 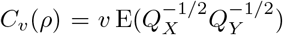 (for *v* > 2)—which depends on *ρ*. Then

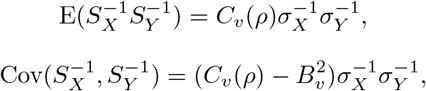

and equation (3) becomes

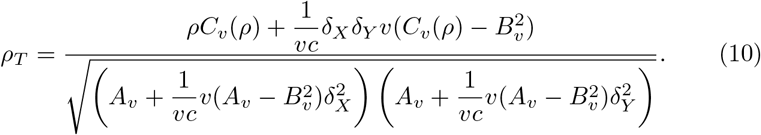

One can show that, under the conditions of Theorem 2, as *v* = *n* − 2 → ∞, 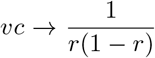, *A*_*v*_ → *B*_*v*_ → 1, and 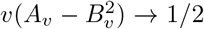, and thus 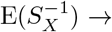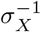 and 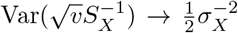. The asymptotic result in Theorem 2 suggest that as *v* → ∞,

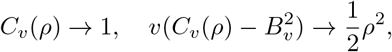

but we did not find a direct analytical proof for these limits.

## Estimating the variance inflation factor

The variance inflation factor (VIF) for a set of m test statistics, *t*_1_, …, *t*_*m*_, is defined as

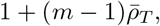

where 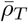 is the average of all pairwise test-statistic correlations (*ρ*_*T*_ ’s). In the case of two-sample *t*-test, given the data-row correlations and mean differences between the two groups in all data rows, we can use equation (4) to compute *ρ*_*T*_ for all row pairs, and in turn the VIF. If we consider the two-sample *t*-test performed on *m* = 21 data rows in a matrix with correlated data rows (*ρ* = 0.1 for all pairs, variance σ^2^ = 1 for all rows) and mean differences ranging from −3 to 3 (uniformly spaced, i.e., *δ* = −3, −2.4, −1.8, …, 3 for the 21 rows) between two groups (*n*_1_ = *n*_2_ = 30), the true VIF value computed using test-statistic correlations should be 2.48; the VIF computed using the data-row correlations is 3.00, which overestimates the true VIF. In practice, for each data row *i* = 1, …, *m*, the mean difference *d*_*i*_ can be estimated by sample mean difference. Let 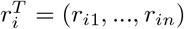 be the vector of residuals after fitting the two group means, then σ_*i*_ can be estimated by 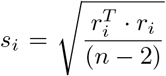. Between a pair of data rows *i* = *X, Y*, the data-row correlation *ρ* can be estimated by the sample correlation coefficient between the residual vectors *r*_*X*_ and *r*_*Y*_, and the test-statistic correlation *ρ*_*T*_ is estimated by plugging in estimated values of *ρ, d*_*X*_, *d*_*Y*_, σ_*X*_, σ_*Y*_ into equation (4). We simulated *H* = 5000 data matrices with the above specified *m, n*_1_, *n*_2_, *ρ*, and *δ* values, and estimated VIF by either using the estimated test-statistic correlations or directly replacing the test-statistic correlations by the estimated data-row correlations. Figure 3 summarizes the histograms of the estimated VIF values, we can see using the estimated *ρ*_*T*_ values to compute the VIF gives less biased results; using the data-row correlations in place of the test-statistic correlations tends to overestimate the VIF.

**Figure 3:**
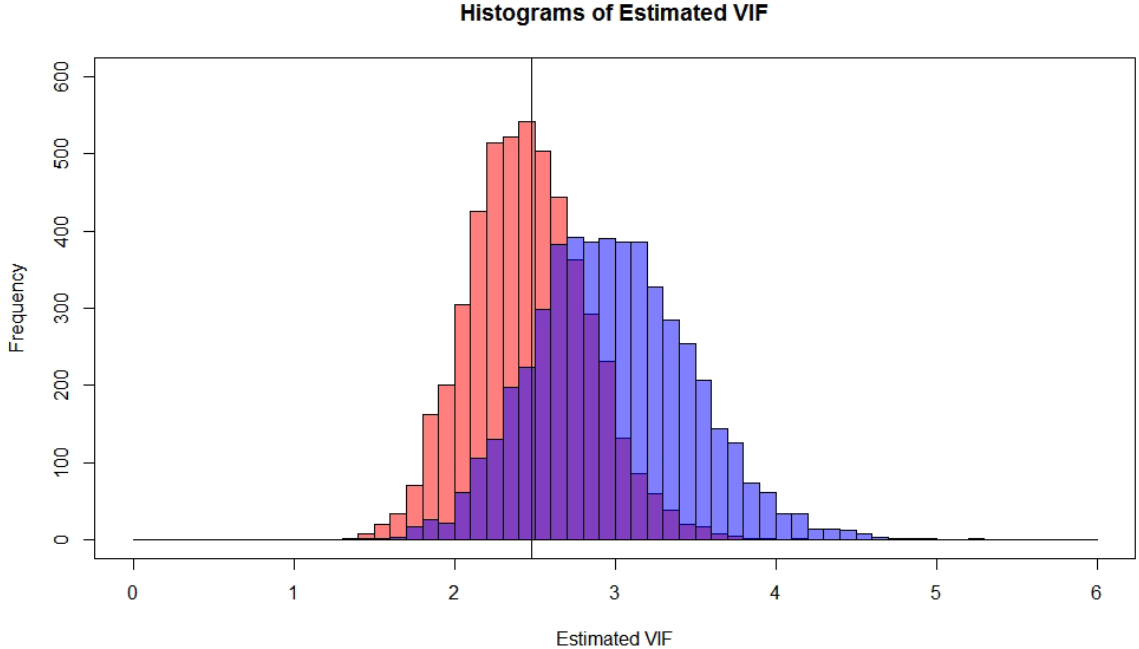
Histograms of the VIF values estimated by estimating *ρ*_*T*_ for all pairs (left) or by replacing *ρ*_*T*_ by corresponding sample data-row correlations (right). The vertical line indicates the true VIF value of 2.48.

## 3.1. Pairwise correlations among moderated t-test statistic values

We use simulation to examine pairwise test-statistic correlations for the moderated *t*-test as implemented in the R package `limma` ([15]) when the data rows are correlated. When computing the moderated *t*-statistic, the variance estimations for the data rows are shrunk towards a common value. Effectively, the variance estimation for each data row draws information from all data rows. This makes deriving an analytical formula for the pairwise test-statistic correlation difficult.

To examine the correlation among test statistic values, we need to simulate multiple data sets. To see the effect of shrinkage, it is also necessary that we simulate many data rows in each data set. In each data set, we simulated 1000 rows of normal data from two samples of size 10 each—resulting in a 1000 × 20 data matrix. In the control group (first 10 columns), the mean was set to 0; in the treatment group (the last 10 columns), the mean was simulated to be −3 for rows 1–50, 2 for rows 51–100, and 0 for rows 101–1000 respectively. The variance was 1 for all data points, the correlation between any pair of data rows was simulated to be *ρ* = 0.1 or *ρ* = 0.5. We simulated *H* = 10000 date sets under each *ρ* value. To fix ideas, one can think each simulated data set as representing normalized (log-transformed) gene expression data from microarrays. The different mean levels under treatment ({−3, 2, 0}), represent different degrees of differential expression (DE). The parameter settings were kept simple here—3 DE classes (*δ*_*X*_, *δ*_*Y*_ ∈ {−3, 2, 0}) and a constant data-row correlation (*ρ* = 0.1 or 0.5)—just enough to reveal the connection between data-row correlations and test-statistic correlations as (*δ*_*X*_, *δ*_*Y*_) vary.

In each data set, the moderated *t*-statistic value was computed for each row. Pairwise sample test-statistic correlations were then computed over the 10000 independently simulated data sets—giving a 1000 × 1000 matrix of sample pairwise test-statistic correlations. We grouped these pairwise test-statistic correlations according to (*δ*_*X*_, *δ*_*Y*_) values and summarized the distribution in each group in Figure 4 (data-row correlation is fixed at *ρ* = 0.1 or *ρ* = 0.5). The main conclusions are:

1. Test-statistic correlations are generally not same as data-row correlations and their relationship depends on the DE statuses (i.e., depending on (*δ*_*X*_, *δ*_*Y*_) values)—this is similar to the standard two-sample *t*-test case that we explored in the paper.
2. However, the relationship between test-statistic correlation and data-row correlation does not follow the analytical formula derived for the two-sample *t*-test case. In particular, we see that when both data rows are from the same DE class (*δ*_*X*_ = *δ*_*Y*_ = −2 or *δ*_*X*_ = *δ*_*Y*_ = 3), the test-statistic correlations tend to be greater than the data-row correlation. For the usual two-sample *t*-test statistic, we have seen earlier that the magnitude of test-statistic correlation tends to be less than that of the corresponding data-row correlation.

**Figure 4:**
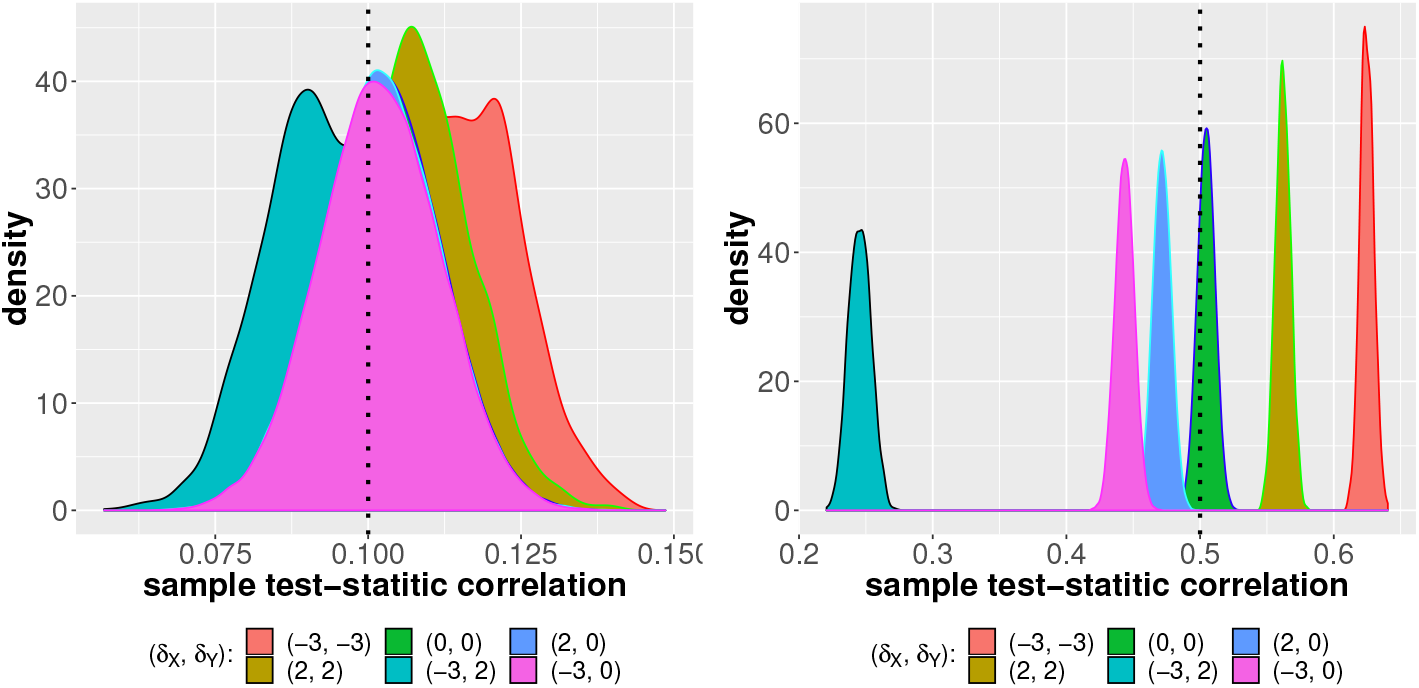
Distribution of pairwise sample test-statistic correlations for the moderated-*t* test. The pair-wise test-statistic correlations are grouped by the (unordered) values of (*δ*_*X*_, *δ*_*Y*_) and the distribution in each group is plotted. The data-row correlation is fixed for all pairs at *ρ* = 0.1 (left panel) or *ρ* = 0.5 (right panel) and is indicated by the vertical line in each plot.

